# Multivariable Mendelian Randomization adjusting for heritable confounding analyzes the causal effects of C-reactive protein on multiple diseases

**DOI:** 10.1101/2025.01.08.631816

**Authors:** Ruoyao Shi, Jean Morrison

## Abstract

**Background:** C-reactive protein (CRP) is a marker of inflammation associated with autoimmune, cardiovascular, and neuropsychiatric disorders. However, it remains unclear whether CRP causally affects these traits or if observed associations result from reverse causation or confounding. Mendelian randomization (MR) uses genetic variants as instrumental variables to estimate causal effects and avoid the biases present in observational studies. Prior MR studies have suggested causal effects of CRP on several traits, including schizophrenia, bipolar disorder, and colorectal cancer. However, MR may produce biased results if factors that confound the exposure and outcome are heritable, resulting in horizontal pleiotropy. This is a major concern for studies of CRP, because CRP levels may increase in response to inflammation caused by a wide range of heritable conditions.

**Methods:** Multivariable Mendelian randomization (MVMR) can be used to eliminate bias from heritable confounding when GWAS summary data are available for confounders. In this study, we use MVMR to estimate the causal effects of CRP on 12 outcomes with prior evidence of a causal or associational link to CRP. We use a novel computational pipeline to identify a broad set of potential heritable confounders between CRP and each outcome trait from studies in the MRC-IEU OpenGWAS database. We compare MVMR results with computationally selected confounders to univariable MR results and MVMR using a narrower, literature derived set of confounders.

**Results:** We find that univariable MR suggests evidence of a potential risk-increasing effect of CRP on coronary artery disease, knee osteoarthritis, and rheumatoid arthritis, and a protective effect on schizophrenia. However, after adjusting for computationally selected heritable confounders, only the causal effects on rheumatoid arthritis (OR 1.18, 95% CI [1.07,1.31], p=0.0010 by GRAPPLE) and schizophrenia (OR 0.87, 95% CI [0.79,0.96], p=0.0038 by GRAPPLE) remain significant. Additionally, after adjusting for confounders we find evidence of a potential protective effect of CRP on bipolar disorder at the nominal significance level, which is not observed in the univariable analysis.

**Conclusion:** These results suggest that univariable MR analyses of CRP may be biased by high levels of heritable confounding, though CRP may indeed play a causal role in development of some diseases, potentially mediated by its role in innate immunity. These results also highlight the potential for automatic confounder selection to improve the robustness of Mendelian randomization analyses.

**Key Messages:**

- We used multivariable Mendelian randomization to estimate the causal effect of CRP levels on various diseases after adjusting for heritable confounders.
- We proposed a novel computational pipeline for phenome-wide heritable confounder selection using traits from the MRC-IEU OpenGWAS database.
- Our study did not find evidence of a causal effect of CRP on multiple diseases, except rheumatoid arthritis and schizophrenia.
- Our study highlights the role of CRP as an indicator of inflammation rather than a causal factor in disease risk, suggesting that previous MR analyses may have been biased by heritable confounding.

## 1 Introduction

Inflammation is a complex biological response triggered by the immune system in response to injury, infection, or tissue damage [1]. While inflammation is a vital protective mechanism, dysregulated or chronic inflammation can contribute to the development and progression of various human diseases [2], such as cardiovascular diseases [3], neuropsychiatric disorders [4, 5], autoimmune diseases [6], and cancers [7].

C-reactive protein (CRP), an acute-phase reactant produced by the liver in response to inflammation, is a commonly used biomarker of inflammation and infection in clinical practice [8]. Many conventional observational studies have reported that CRP level is associated with cardiovascular diseases, such as coronary artery disease and stroke [9], neurological diseases such as Alzheimer’s disease [10] and schizophrenia [11], type 2 diabetes [12], rheumatoid arthritis [13], inflammatory bowel disease [14], and some cancers [15]. However, these associations do not necessarily indicate a causal effect of increased CRP level on risk of these diseases. Observational associations may also occur if there are reverse effects of disease status on CRP level or confounding factors that are common causes of both increased CRP level and disease risk.

Mendelian randomization (MR) [16] is a form of instrumental variable analysis that uses genetic variants, typically single-nucleotide polymorphisms (SNPs), as instruments to estimate the causal effect of an exposure on an outcome. MR offers several advantages over observational studies, including robustness to reverse causality and unmeasured confounding. However, one major issue in MR studies is heritable confounding, which occurs when genetic instruments affect a common cause of the exposure and outcome, leading to bias in the MR estimates [17, 18]. Many potentially confounding traits, including behavioral traits and biomarker levels, have been shown to be heritable in recent years, highlighting the importance of this issue for MR studies. Robust univariable MR methods have been developed to account for heritable confounding [18, 19, 20, 17, 21]. However, these methods generally rely on an assumption that a majority or a plurality of instruments are not mediated by heritable confounders. Alternatively, if confounders are known and GWAS associations are available, confounders can be adjusted for using multivariable Mendelian randomization (MVMR) [22, 23, 24, 25].

Previous MR studies have identified evidence of a causal effect of CRP on several diseases and traits, including schizophrenia, inflammatory bowel disease, and rheumatoid arthritis (Table 1). However, these studies have either used univariable MR without incorporating the effect of any heritable confounders or adjusted only for body mass index (BMI). Given CRP’s role as a nonspecific marker of inflammation, its levels may be influenced by a range of genetic variants associated with both inflammation and diseases. This means that MR studies of CRP could be particularly susceptible to heritable confounding. Consequently, prior MR findings may suffer from residual heritable confounding, which could introduce bias to the estimates.

**Table 1:**
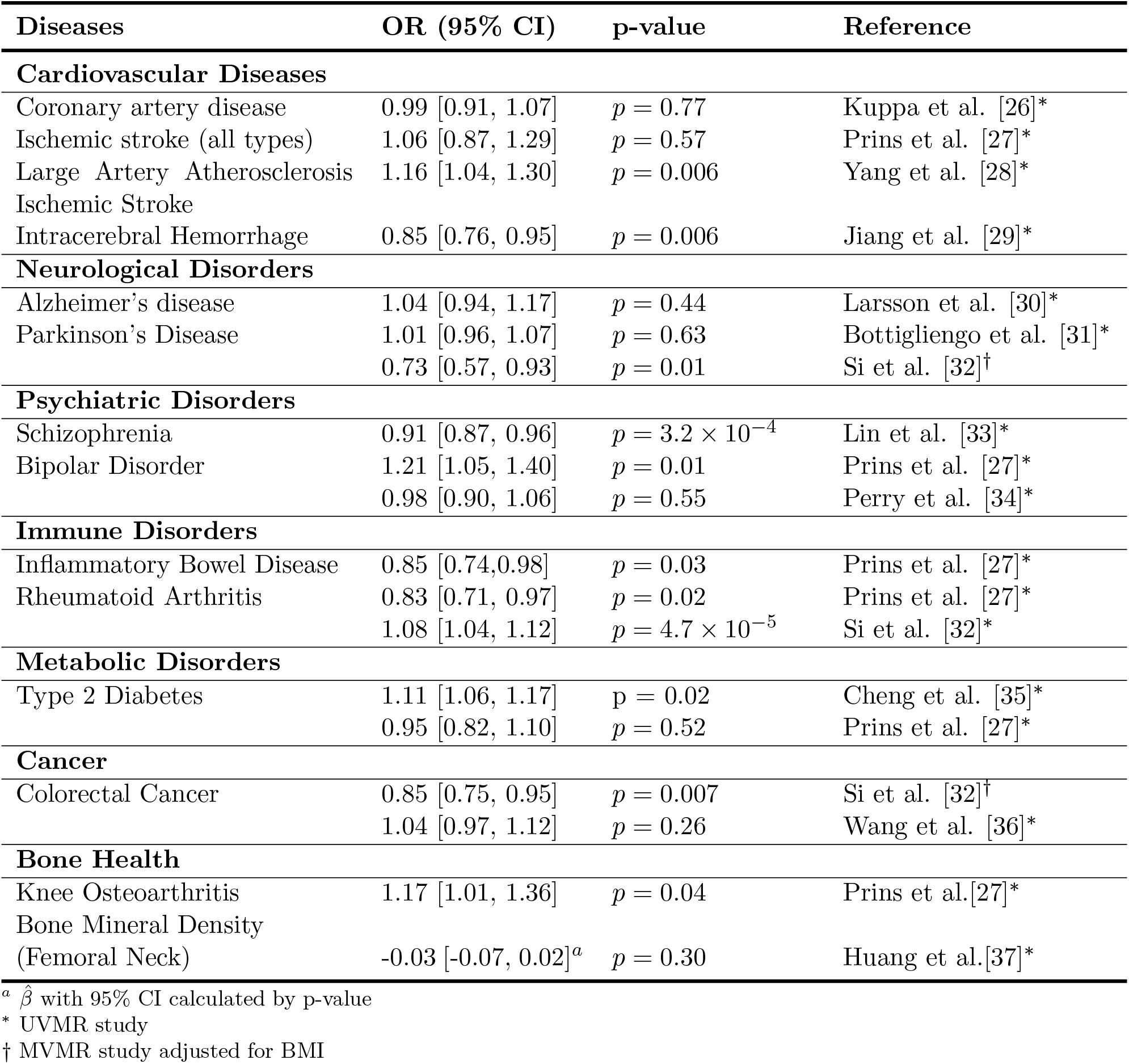
Results summary between CRP-level and diseases in previous studies.

BMI, smoking status, comorbidities such as diabetes, and physical activity are among the classical confounders commonly accounted for in previous CRP research [38, 39, 40, 41, 42]. However, these may not encompass all necessary variables.

In this study, we perform a comprehensive re-analysis of the causal effects of CRP on 12 outcome traits using a large-scale computational pipeline to identify potential heritable confounders among traits present in the MRC-IEU OpenGWAS Database. We find that adjusting for confounders eliminates evidence of a direct causal effect of CRP for all but two of the 12 outcomes. These results are likely not due to weak instrument bias, as the F-statistics remain strong post-adjustment. Additionally, our sensitivity analyses confirm the robustness of these findings across different criteria for confounder selection process. Evidence of a causal effect of CRP persists after confounder adjustment for two traits, schizophrenia and rheumatoid arthritis. These results may indicate a causal role of immune response in the etiology of these two diseases, though further validation is required to verify these effects.

## 2 Material and Methods

### 2.1 Data description and genetic instruments of CRP

We obtained publicly available GWAS summary statistics from MRC Integrative Epidemiology Unit (IEU) OpenGWAS database for CRP-level and 12 disease outcomes, schizophrenia (SCZ), bipolar disorder (BD), Alzheimeri’s disease (AD), Parkinson’s disease (PD), stroke, coronary artery disease (CAD), type 2 diabetes (T2D), colorectal cancer (CRC), knee osteoarthritis (OA), inflammatory bowel disease (IBD), rheumatoid arthritis (RA), and bone mineral density (BMD). We obtained genetic associations for CRP from a recent European ancestry GWAS of approximately 575k individuals [43]. All GWAS for outcome traits were also performed in European ancestry cohorts. The specific description of all GWAS studies is listed in Supplementary Table S1.

We selected variants associated with CRP that exceeded a genome-wide significance threshold of *p <* 5 *×* 10^*−*8^. We performed linkage disequilibrium (LD) clumping with a 10,000 kb clumping window and a threshold of *R*^2^ *<* 0.001, using the European LD reference panel from the 1000 Genomes Project data, to ensure the independence of genetic associations. Ultimately, we identified 264 SNPs as genetic instruments for CRP. We were not able to select instruments in an external data set because all large European ancestry studies of CRP level have substantial sample overlap with our primary data source. Performing in-sample instrument selection may induce a small amount of bias in causal estimates due to winner’s curse, however, this is a common practice in MR.

### 2.2 Phenome-wide identification of candidate confounders

We developed a comprehensive pipeline to identify potentially confounding traits present in the MRC-IEU OpenGWAS database, described in Figure 1.

**Figure 1:**
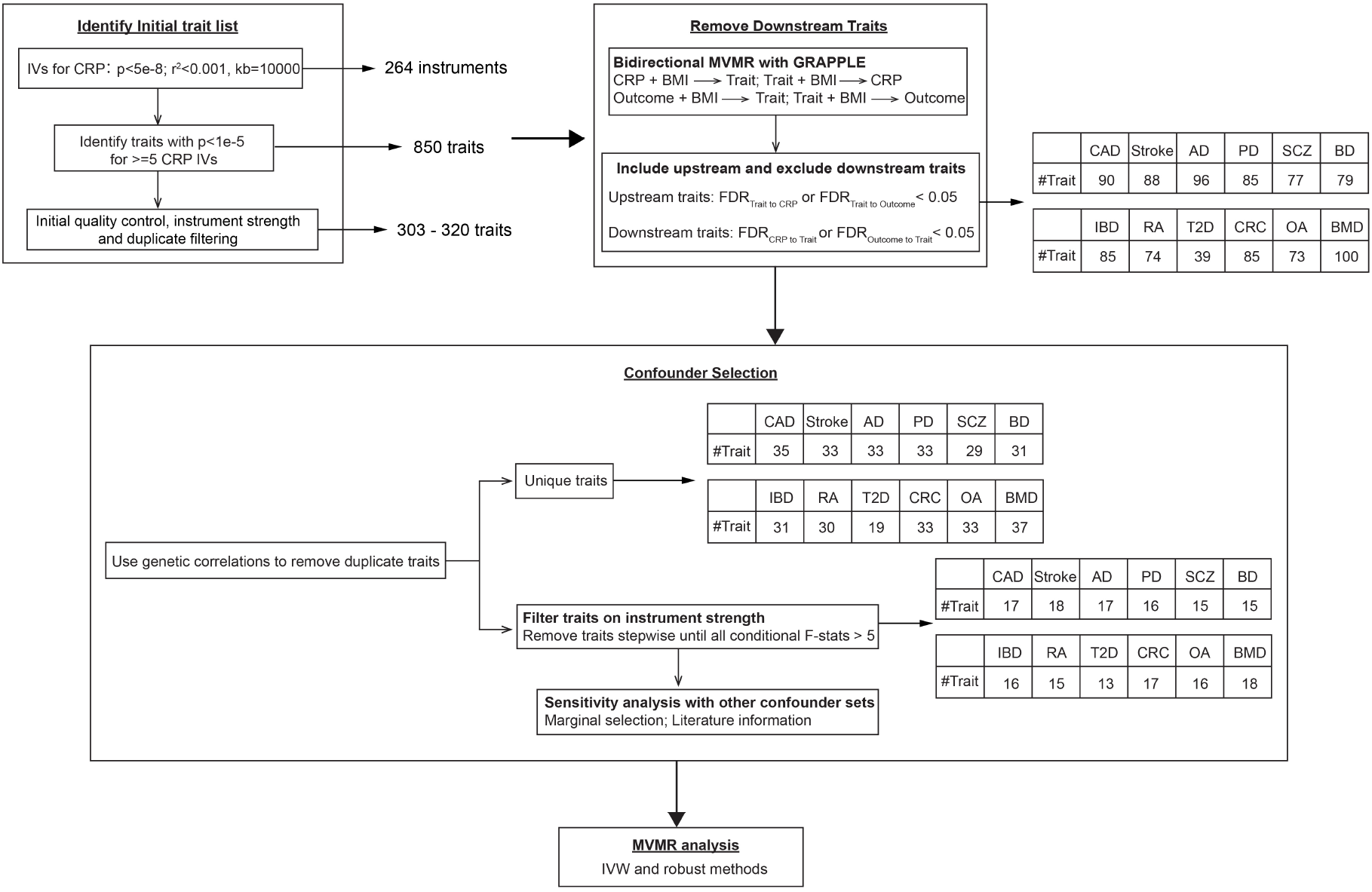
Workflow of heritable confounder selection.

#### 2.2.1 Initial trait identification and quality control

Our initial trait identification step is intended to permissively identify traits that could be bias-inducing heritable confounders of CRP and outcome traits from a set of 5,746 studies that make up four batches of the MRC-IEU OpenGWAS database (ukb-b: IEU analysis of UK Biobank phenotypes, ieu-a and ieu-b: GWAS summary datasets generated by various consortia and manually curated, and ebi-a: datasets imported from the EBI database of complete GWAS summary data). Since confounding traits must be causal for CRP, we first identified traits associated with at least 5 of the 264 CRP instruments at a relaxed threshold of *p <* 10^*−*5^, which identified 850 traits initially. We retained studies that met this criterion, were conducted in European ancestry populations, were not sex-specific, and had more than 1 million variants.

To quickly group together duplicate traits, we performed greedy selection based on the Jaro-Winkler string similarity of trait names [44]. We first ranked traits according to the number of genome-wide significant variants after LD clumping (*p <* 5 *×* 10^*−*8^; clumping distance cutoff = 10,000 kb, clumping *R*^2^ cutoff = 0.001). We then iteratively selected the top trait and removed all traits with a similarity score greater than 0.95 with the selected trait. This procedure continued until all traits were either selected or removed. This allowed us to remove studies for traits with exactly or nearly exactly the same name as a selected study.

Finally, we performed initial filtering to remove traits with only weak instruments. For each candidate confounder, we computed the conditional F-statistic [23] for the model including CRP, BMI, and the candidate confounder as exposures for each outcome. If the conditional F-statistic for the candidate confounder was below 5 for a given outcome, we eliminated that trait as a candidate for that outcome. We note that the conditional F-statistics may differ slightly across outcomes due to differences in the variants available in each outcome study. After these steps, there were 303-320 studies left for different outcomes.

#### 2.2.2 Removal of Causally Downstream Traits

Heritable confounders are causally upstream of both the exposure and outcome. Traits which are causally downstream of either the primary exposure or the outcome should not be included in the MVMR analysis. Downstream traits do not improve the estimate of the effect of the primary exposure on the outcome, and can substantially reduce precision, or even introduce bias [23]. Therefore, we used bidirectional MR to identify and remove downstream traits. Given that BMI has a strong and established effect on CRP [45, 46] and may plausibly affect many of the outcome traits, we selected BMI as a prior known confounder and included it in the bidirectional MR analysis. Additionally, to assess the impact of adjusting for BMI, we conducted a parallel set of analyses without adjusting for BMI in the bidirectional MR step, or using BMI in initial instrument strength filtering step.

For each potential confounder-outcome combination, we performed two MVMR analyses using GRAPPLE [24]. In one analysis, we estimated the direct effects of the potential confounder and BMI on the outcome, and in the other, we estimated the direct effects of the outcome and BMI on the potential confounder. Additionally, for each potential confounder, we used MVMR to estimate the effects of the potential confounder and BMI on CRP level as well as the effects of CRP level and BMI on the potential confounder.

In each MVMR analysis, we removed very large effect variants, defined as SNPs with an absolute standardized effect size (SD/SD scale) greater than 0.1 for all exposures. We implemented this step because very high-effect variants may be related to full-body syndromes increasing the risk of pleiotropy. These variants also tend to have a large effect on estimates, due to being high leverage points. For example, variants in the APOE locus have a very strong effect on AD risk [47], as well as effects on lipids and other biomarkers [48]. This locus is generally understood to be highly pleiotropic and is often removed in MR studies of AD [49, 50].

We removed an average 0.67 high-effect SNPs per analysis. Following this, we applied Steiger filtering [51] to remove variants more strongly associated with the outcome than with the exposure. For simplicity in this step, we treated all potential confounders as continuous traits, while disease outcomes were treated as binary traits, incorporating their respective disease prevalence, given in Supplementary Table S1. The final set of instruments comprised the union of the selected SNPs from all exposures.

We applied the Benjamini-Hochberg procedure to estimate the false discovery rate (FDR) from *p*-values obtained from our bidirectional MVMR analysis separately for each outcome and CRP level. We retained traits with an estimated FDR below 0.05 for their effect on either CRP level or the outcome, and excluded any with an estimated FDR below 0.05 for effects from CRP level or outcome. If a trait met both criteria, it was excluded. This selection process resulted in a refined potential confounder list of 39-100 traits for each outcome.

As a sensitivity analysis, we repeated this selection process using an FDR threshold of 0.1 rather than 0.05. We found that the selected confounders remained largely consistent. Specifically, 84% of traits selected using the 0.05 threshold are also selected at the 0.1 threshold. Additionally, the final causal estimates were not sensitive to the difference in selected traits (Supplementary Table S2-13).

#### 2.2.3 Removal of Duplicated Traits

To eliminate highly similar or duplicated traits that were not clumped together by the Jaro-Winkler score previously, we performed a similar greedy selection procedure, this time using genetic correlation as the measure of trait similarity, and a threshold of 0.8. As before, traits were ranked by the number of genome-wide significant variants. We additionally chose to favor continuous traits over binary traits. If a binary trait was selected and there existed a continuous trait with genetic correlation greater than 0.8, we would select the continuous trait instead. Additionally, we removed traits related to medication usage in this step because these traits are likely to be causally downstream of disease status. There were 19-37 traits selected for each outcome after this step.

#### 2.2.4 Instrument Strength Filtering

Traditional MVMR estimation using inverse variance weighted regression can be biased by inclusion of too many weak instruments. To mitigate this risk, we have based our analysis on the GRAPPLE [24] and MRBEE [25] methods, which are more robust to weak instruments. However, even using a weak instrument robust method, it is still necessary to remove traits with only weak genetic associations to avoid identifiability issues.

We used a stepwise approach to remove traits with weak instruments. We first computed the conditional F-statistic [23] for all candidate confounders. If the lowest F-statistic was below 5, we removed the trait with the lowest F-statistic. We repeated this process until all remaining traits had a conditional F-statistic greater than 5. This resulted in 13-18 potential confounding traits selected per outcome. Without filtering traits based on instrument strength, we observed that the causal estimates for some traits could have very large standard errors.

### 2.3 Alternative Trait Selection Methods

In our primary analysis, we adjusted for all confounders remaining after stepwise instrument strength filtering. As a sensitivity analysis, we also considered a more aggressive, marginal selection criterion in which traits were only included if our bidirectional MVMR analysis indicated evidence of an effect on both CRP level and the outcome at the FDR *<* 0.05 level. Moreover, We also considered augmenting our selected list of confounders with three traits, smoking, diabetes, and physical activity, which were identified based on previous information that these traits may affect CRP level [38, 39, 41, 42]. Finally, we performed a sensitivity analysis adjusting only for BMI, which is the most common approach to MR analysis of CRP after unadjusted univariable estimation.

### 2.4 Mendelian Randomization Analysis

We performed multivariable MR analysis to estimate the causal effect of CRP-level on the 12 outcome traits, adjusting for selected confounders. We selected instruments using the same procedure described for the bidirectional screening step used to identify causally downstream traits. We first selected genome-wide significant variants (*p <* 5*×*10^*−*8^) for CRP level and each confounder. We then performed LD clumping, removed variants with standardized effects greater than 0.1 (average 3.8 variants removed per analysis), and finally performed Steiger filtering.

We used GRAPPLE [24] as our primary analysis method. Supplementary Tables include results from the MRBEE [25] and MV-IVW methods [52]. Both GRAPPLE and MRBEE can adjust for sample overlap using a residual correlation matrix, which we estimated using cross-trait Linkage Disequilibrium Score Regression (LDSC). We note, however, that MV-IVW results may not be reliable when adjusting for a large number of confounders.

For comparison, we computed univariable MR estimates for the effect of CRP-level on each outcome using the same methods and instrument selection procedures.

## 3 Results

### 3.1 Summary of Selected Potential Confounders

We identified some traits with evidence of causal effects on CRP conditional on BMI, including phosphate levels, peak expiratory flow, ankle spacing width, urea levels, triglycerides, basophil percentage of white cells, and diastolic blood pressure. These were consistently included in analyses across multiple outcomes.

Additionally, we identified several outcome-specific traits, including testosterone levels and frequency of childhood sunburn for BMD; serum uric acid levels for RA; hypothyroidism or myxedema and serum creatinine levels for T2D; glucose levels and mean platelet volume for CAD. The complete list of the final selected confounders is provided in Supplementary Table S2-13. On average, we selected approximately 16 confounders for each outcome.

### 3.2 MR Analysis Results

UVMR results suggest evidence of causal effects of CRP level on OA, CAD, SCZ, and RA at the nominal *p <* 0.05 level. However, after adjusting for computationally selected heritable confounders, we found that only effects on RA and SCZ remain significant (Figure 2). Additionally, in the adjusted analysis, the effect of CRP on BD is nominally significant, while it was not in the UVMR analysis.

**Figure 2:**
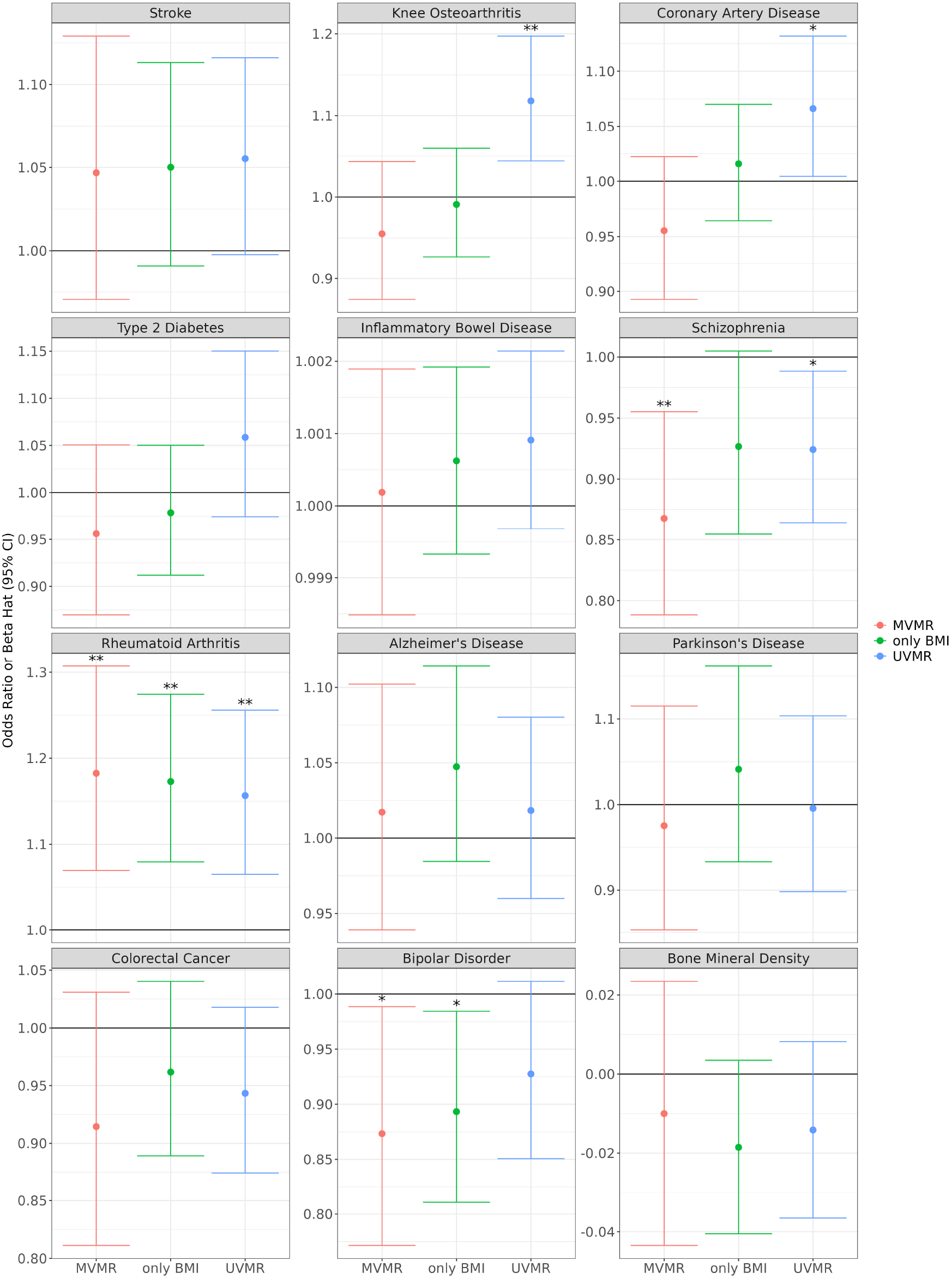
Estimated causal effect of CRP-level on disease outcomes estimated using GRAPPLE. Effect estimates (points) and 95% confidence intervals are shown. Asterisks indicate significance level (* *p <* 0.05; ** *p <* 0.05*/*12) The MVMR analysis included all computationally selected candidate confounders. Only BMI was included in the “only BMI” analysis.

Both UVMR and MVMR analysis indicate a protective causal effect of CRP on the risk of SCZ. Estimates of the effect of CRP on RA were positive in both UVMR and MVMR analysis.

Employing either UVMR or MVMR analysis adjusting for selected confounders, no significant causal relationships were detected between genetically determined plasma CRP levels and the risk of T2D, IBD, AD, PD, CRC, and BMD. We note that univariable IVW does identify a causal effect of CRP on stroke and IBD, though this method is more sensitive to pleiotropy than GRAPPLE (Supplementary Figure S1).

We compared our UVMR-IVW results with findings from previous studies. Both our analysis and prior research found no significant causal relationships between CRP and AD, CAD, BMD, and consistent significant causal effects for OA, SCZ, and RA. However, our results revealed some inconsistencies for PD, BD, CRC and T2D, where previous studies have shown mixed results, some reporting significant causal effects while others found null results. These discrepancies could stem from variations in study designs, sample size, and genetic instruments selected by different studies.

### 3.3 Effects of BMI Adjustment

Adjusting for BMI in the bidirectional MR step impacted the set of selected traits. Traits such as triglycerides, phosphate levels, and urea levels were consistently identified, whether BMI was adjusted for or not. In sensitivity analysis not adjusting for BMI, we additionally identified several liver enzyme-related traits, including alanine transaminase, aspartate aminotransferase levels, and insulin-like growth factor-I as potential confounders. These traits were not selected in the BMI adjusted analysis because, adjusting for BMI, we identify a significant effect of CRP on these traits, meaning that they may be downstream traits of CRP. Additionally, adjusting for BMI enabled the identification of potentially important traits such as diastolic blood pressure and 25 hydroxyvitamin D levels that were not identified in the unadjusted sensitivity analysis. These findings underscore the potential impact of BMI as a confounder in MR analyses of CRP and support the inclusion of BMI in the bidirectional MR step. The results without BMI adjustment are provided in the Supplementary Table S2-13.

MVMR analysis adjusting only for BMI generally yielded the same conclusions as our MVMR analysis, which adjusted for a broader set of selected confounders (Figure 2, Supplementary Figure S1, Supplementary Figure S2). This consistency underscores the significant confounding effect of BMI in these relationships. However, this was not true in all cases. For example, adjusting only for BMI resulted in a null causal effect between CRP and SCZ while adjusting for selected confounders indicated a significant positive relationship. Adjusting for a comprehensive set of selected confounders also resulted in more consistent conclusions across MVMR methods. This could occur if confounder adjustment results in a decrease in heterogeneity, which is modeled differently by the different methods. This suggests that, while BMI is an important confounder, our thorough confounder selection process can enhance the reliability of causal estimates by identifying a more complete list of potential confounders.

## 4 Discussion

In this study, we investigated whether genetically determined CRP levels are causally associated with the risk of multiple human diseases after adjusting for potential confounders using MVMR. Our findings suggest that previous UVMR analyses of CRP may have produced biased causal estimates due to heritable confounding, and CRP may have fewer true causal effects on human diseases than previously reported. Additionally, in some cases such as BD, heritable confounding may obscure causal effects. Furthermore, we introduced a comprehensive pipeline to identify heritable confounders, generating a complete confounder list, and thereby enhancing the robustness of causal inferences obtained through MVMR.

Our analysis identified BMI as a significant source of heritable confounding, along with other important confounders such as apolipoprotein B levels, diastolic blood pressure, waist-hip ratio, and triglycerides. Findings from MVMR analysis indicate that CRP may primarily act as a biomarker of inflammatory responses and related conditions, rather than having a direct causal impact on the risk of human diseases.

We identified a potential positive causal relationship between CRP levels and the risk of RA after adjusting for selected confounders. This finding aligns with preclinical evidence suggesting that CRP is involved in the pathogenesis of RA [53], particularly in bone destruction processes. CRP has been shown to induce the expression of receptor activator of nuclear factor-*κ*B ligand (RANKL) [54], thereby stimulating osteoclastogenesis and bone resorption, which are crucial processes in the progression of RA. In concordance with the established role of CRP as a biomarker for RA disease activity, our findings suggest that CRP might also play a contributory role in the disease’s development. Further research is needed to fully understand the complex biological role of CRP in RA progression and its implications for treatment.

In our study, we observed that genetically elevated CRP levels were associated with a protective causal effect on the risk of developing schizophrenia and bipolar disorder, after adjusting for confounding factors, which aligns with previous MR study findings [27]. However, this finding contrasts with the elevated CRP levels frequently observed in schizophrenia patients, as reported by previous observational studies [55, 56, 57]. This discrepancy suggests that elevated CRP levels in these patients may reflect a secondary inflammatory response, potentially triggered by the disorder itself or environmental factors. Our results indicate that genetically elevated CRP might signify a more robust immune defense, which could provide protection against infections or immune challenges speculated to contribute to the development of schizophrenia [58, 59].

Several limitations exist for this work. First, our confounder search was limited to traits available within selected batches of the MRC-IEU OpenGWAS database, and sex-specific or non-European studies were also not considered in this study. Therefore, the confounder list may be incomplete. Second, in the bidirectional MVMR step, only accounting for BMI may not be sufficient to provide fully accurate estimates. Third, although we employed multiple robust MVMR methods to mitigate biases arising from weak instruments, horizontal pleiotropy, and measurement errors, these issues cannot be entirely eradicated. Fourth, potential bias could be introduced by instrument selection of the specific CRP study. Lastly, our MR analyses reflect the genetic component of lifelong exposure to altered CRP levels. However, it is possible that CRP levels during specific life stages, such as early development, might have significant effects on disease risk. We obtained estimates for the SNP-CRP associations from adult subjects, but these associations might differ at other time points in life.

