## Supplementary Table S1, Supplementary Figure S1, Supplementary Figure S2 for "Multivariable Mendelian Randomization adjusting for heritable confounding analyzes the causal effects of C-reactive protein on multiple diseases"

### Supplementary Materials

| GWAS ID | Trait | Sample Size | nSNPs | nCase | nControl | PMID | Prevalence |
|---|---|---|---|---|---|---|---|
| ebi-a-GCST90029070 | C-reactive protein levels | 575,531 | 10,713,245 | NA | NA | 35459240 | NA |
| ieu-b-5102 | Schizophrenia | 127,906 | NA | 52,017 | 75,889 | 35396580 | 0.0045 [1] |
| ieu-b-41 | Bipolar Disorder | 51,710 | 13,413,244 | 20,352 | 31,358 | 31043756 | 0.0049 [2] |
| ebi-a-GCST90027158 | Alzheimer's disease | 487,511 | 20,921,626 | 39,106 | 46,828 | 35379992 | 0.0068 [3] |
| ieu-b-7 | Parkinson's disease | 482,730 | 17,891,936 | 33,674 | 449,056 | NA | 0.00151 [4] |
| ebi-a-GCST005838 | Stroke | 446,696 | 7,633,440 | 40,585 | 406,111 | 29531354 | 0.0118 [5] |
| ebi-a-GCST005194 | Coronary artery disease | 296,525 | 7,904,237 | 34,541 | 261,984 | 29212778 | 0.05 [6] |
| ebi-a-GCST006867 | Type 2 diabetes | 655,666 | 5,030,727 | 61,714 | 1,178 | 30054458 | 0.0628 [7] |
| ebi-a-GCST90018808 | Colorectal cancer | 470,002 | 24,182,361 | 6,581 | 463,421 | 34594039 | 0.0015 [8] |
| ebi-a-GCST007090 | Knee osteoarthritis | 403,124 | 29,999,696 | 24,955 | 378,169 | 30664745 | 0.0431 [9] |
| ebi-a-GCST90038683 | Inflammatory bowel disease | 484,598 | 9,587,836 | 4,101 | 480,497 | 33959723 | 0.000843 [10] |
| ebi-a-GCST90018910 | Rheumatoid arthritis | 417,256 | 24,175,266 | 8,255 | 409,001 | 34594039 | 0.0046 [11] |
| ebi-a-GCST90014022 | Bone mineral density | 365,403 | 10,783,906 | NA | NA | 34017140 | NA |

Table S1: Data Description of GWAS Summary Statistics

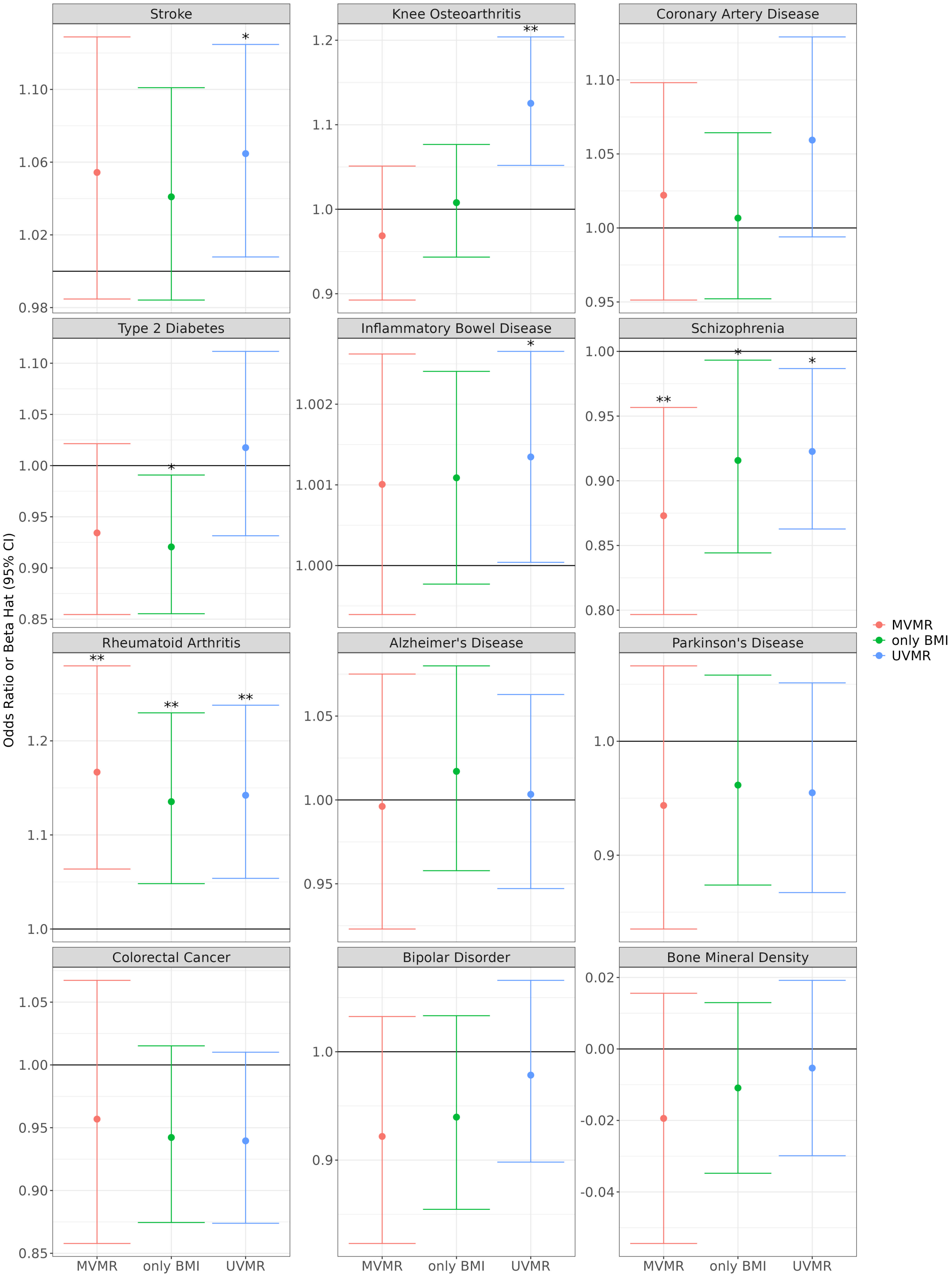


Figure S1: Estimated causal effect of CRP-level on disease outcomes estimated using MV-IVW. Effect estimates (points) and 95% confidence intervals are shown. Asterisks indicate significance level ( * $p < 0.05$; ** $p < 0.05/12$) The MVMR analysis included all computationally selected candidate confounders. Only BMI was included in the "only BMI" analysis.

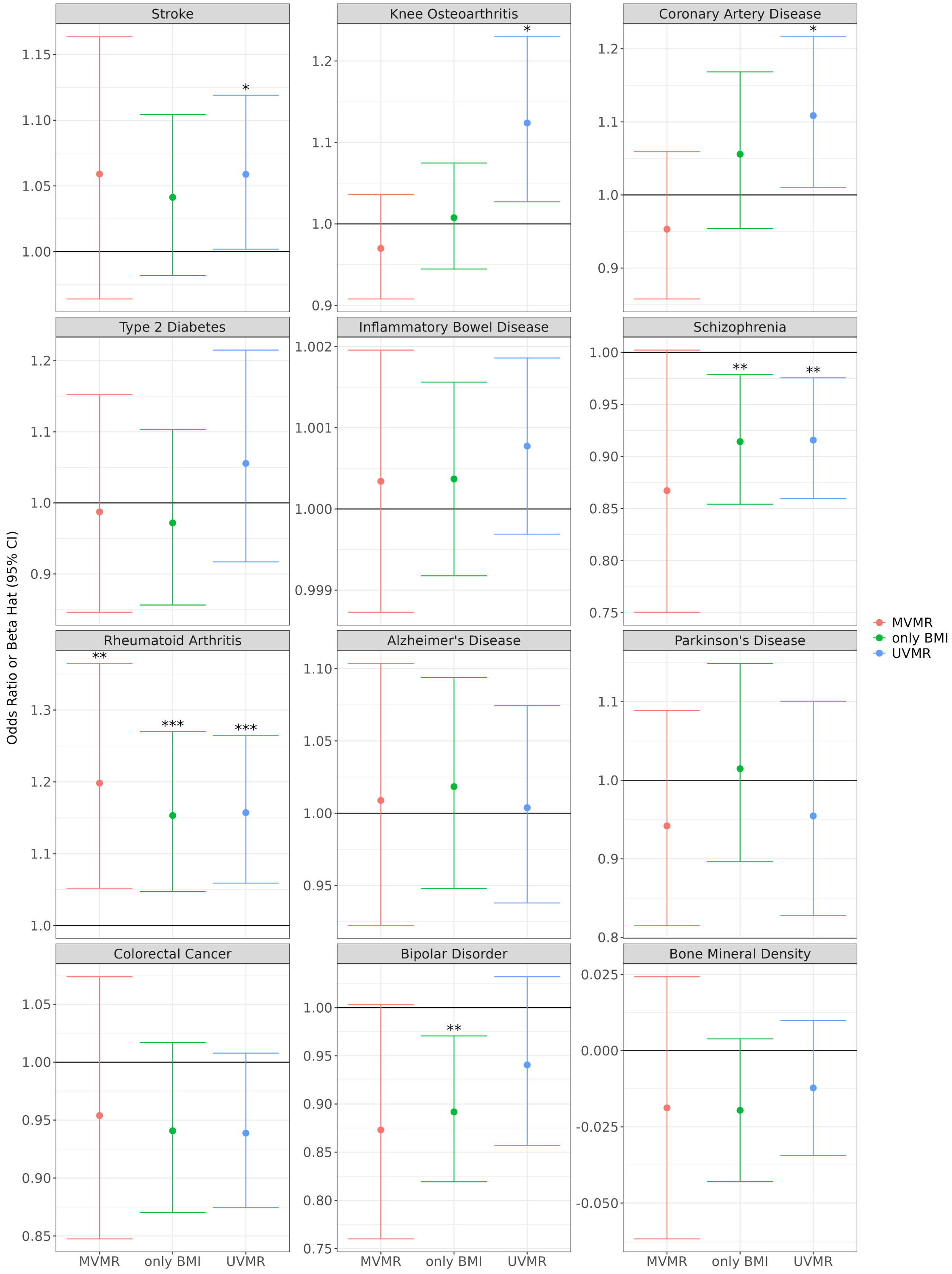


Figure S2: Estimated causal effect of CRP-level on disease outcomes estimated using MRBEE. Effect estimates (points) and 95% confidence intervals are shown. Asterisks indicate significance level ( * $p < 0.05$; ** $p < 0.05/12$) The MVMR analysis included all computationally selected candidate confounders. Only BMI was included in the "only BMI" analysis.
